# Development of a Novel Microfluidic Co-culture model to study Organoid Vascularization

**DOI:** 10.1101/2022.03.25.485813

**Authors:** Lydia S. Schulla, E. Diana Alupoaie, Leanne De Silva, Debby Gawlitta, Sabine Middendorp, Paul J. Coffer, M. Guy Roukens

## Abstract

Organoids hold great potential for regenerative medicine and biomedical research. While they are able to recapitulate many structural and functional aspects of their *in vivo* counterparts, they lack a functional vasculature. In this study, we aimed to coculture human small intestinal organoids (hSIO) with a vasculature in a triple co-culture system with endothelial cells (ECs) and fibroblasts in a microfluidic device. Organoids and micro-vessels favour distinct matrix and medium conditions, which were optimized to sustain growth of all cell types. In addition, we found that organoids exhibit poor survival in monoculture in the microfluidic devices. Interestingly, co-culturing hSIO together with ECs and fibroblasts enhances stemness and survival of hSIO. This effect can be further enhanced by adding the GSK-3β inhibitor CHIR99021 to the medium to enhance Wnt signalling. Direct contact of fibroblasts is not required for either vessel formation or for the effect on stem cell maintenance of organoids, as culturing fibroblasts in side-channels yields larger micro-vessels and better organoid survival. Additionally, we observed that the response of cells to different culture media is strongly dependent on the culture set-up as the second setup allows organoids to be cultured in angiogenic medium. Our results suggest that angiocrine signalling by the vasculature affects stem cell maintenance, which should be further investigated. Our co-culture system offers a platform to study angiocrine signalling in the small intestine, which could in the future be used to unravel different mechanisms behind endothelial mediated intestinal homeostasis and (patho) physiology.

## Introduction

Since their discovery in 2009^1^, organoids have become a fundamental technology that is widely used in biomedical science. Organoids are 3D structures that are derived from stem cells and consists of organ-specific cells that self-organize to recapitulate aspects of the native tissue architecture and function *in vitro*. Traditional 2D cell lines have long been used in biomedical research, but their applicability and relevance is hampered as they are difficult to establish from non-tumorigenic origins and they often lack the presence of differentiated cells. Organoids are derived from adult stem cells (ASC) or (induced) pluripotent stem cells (IPSC) and overcome many of these issues; hence, protocols have been established to derive organoids from a large variety of tissues consisting of a variety of stem- and differentiated cells ^2^.They can be expanded indefinitely, remain genetically stable in culture and can be established from individual biopsies. Therefore, organoids form excellent cellular tools to study development and disease in a highly individualized fashion ^2–4^. The archetype model system for adult stem cell based organoids is the intestine, since it was the first organoid model to be developed^1^. Intestinal organoids have been used to interrogate stem cell maintenance, differentiation processes and regeneration^2^. As a prime example of their ability to model pathophysiological processes, it has been shown that intestinal organoids can serve as a powerful model to predict drug responses for cystic fibrosis ^5^.In addition, tumors are highly amenable to derive patient-specific organoid lines, which is key to develop a personalized medicine approach to cancer. Indeed, concerted efforts have been made to use organoids as patient-specific models for drug testing in cancer, where the tumor organoids are often stored in biobanks for future references^6^. Recent years have seen a growing interest in combining ASC-based organoids with other cell types, such as fibroblast^7^, immune cells^8,9^ or even with microbes^10^, to generate more complex model systems. Finally, organoids have been proposed to be a promising source for cellular replacement or repair in the field of regenerative medicine^2,11^. Various studies have highlighted this potential by transplanting intestinal organoids in mouse models for acute colitis. Intestinal organoids were shown to repopulate a damaged intestinal epithelium, giving rise to all differentiated cells and restoring functionality of the colon^12–14^. These studies open therapeutic avenues for patients suffering from inflammatory bowel disease or with acute intestinal failure. However, to develop replacement organs for humans larger tissue constructs need to be engineered but the major limitation is vascularization^15,16^.

In native human tissues nearly all cells are found within a distance of 100-150 um of a blood vessel that supplies the cells with oxygen and nutrients and removes waste ^17^. In *engineered* tissues and in organoids the lack of a vasculature has been a major constraint as oxygen and nutrients can only be transported by diffusion. This restricts the maximum thickness of organoids to 150-200 um and limits their potential for replacing or repairing organs^11^. As a model system, organoids could also be even more valuable as integrating a blood vasculature system could contribute to multicellular models that better mimic the complex interplay between various cell types in organogenesis, in organ homeostasis and in disease. This is particularly relevant in complex multi-factorial diseases such as inflammatory bowel disease and cancer. Moreover, the blood vessels also have a *paracrine* function via ‘angiocrine’ signaling, by which soluble or membrane bound factors of vascular endothelial cells affect the development and homeostasis of organs in health and disease^17,18^. This signaling is different for each organ which has its own specialized capillary bed that expresses specific factors to support homeostasis of that organ^17,19^. Crucially, angiocrine signaling has an instrumental role in regeneration^20^. In mouse models lung regeneration after pneumonectomy and hepatocyte proliferation after partial liver resection were shown to critically depend on angiocrine signaling^21,22^. Thus, ensuring that engineered tissues can be vascularized upon transplantation is crucial for organ repair/regeneration. A promising strategy to improve this is to pre-vascularize engineered tissues *in vitro*. For skeletal muscle cells and hepatocytes this facilitates the ingrowth of host blood vessels in transplanted grafts^23,24^. Similarly, prevascularization *in vitro* of organoids could be beneficial for their regenerative capacity.

In the past decade extensive efforts ^25–27^ have attempted to generate vascularized tissue constructs. Despite advances in *in vitro* vascularization approaches, thus far the only way to generate fully functional and perfusable vasculature in organoids has been by *in vivo* transplantation of organoids. This has been shown for example by transplanting kidney organoids under the renal capsule of mice and this resulted in vascularization of the transplanted organoid tissue by functional host vessels. Interestingly vascularization also led to significant maturation of the transplanted organoids compared to those in vitro^28^. Various other studies have shown similar results for liver^24^ and brain organoids ^29–31^. However, the field is in need of vascularization of organoids *in vitro*, as this would be a much more easy approach than *in vivo* transplantation. However, vascularization of organoids *in vitro* has proven to be more challenging approach^26^. A promising methodology for such a model would be to use microfluidic platforms for vascularizing organoids. Microfluidics has the advantages of compartmentatlization and high surface area to volume ratio, which is advantageous for developing self-organizing microvessels from endothelial cells ^26^. Indeed Nashimoto *et al* used a device wherein they cultured fibroblast spheroids in a center matrix, which attracted endothelial sprouts which emanated from two side channels^32^. Other groups have used fibroblasts and endothelial cells to self-organize into a vascular network using microfluidic devices. Sobrino *et al* also cultured tumor cells within the same channel that allowed performing drug screens for chemotherapy in combination with anti-angiogenesis therapy^33^. These studies showed the potential of using microfluidic chips for vascularization, but the vascularization of fibroblast spheroids and tumor cell lines are less representative models than (tumor) organoids. In addition, the generation of such microfluidic devices requires the use of specific technology and highly specialized expertise that is not available routinely for most laboratories.

Therefore, here we aimed to develop a model to vascularize *organoids in vitro*. We used a commercially available microfluidic device from AIM Biotech as it is versatile in its use, robust and easy to handle. We developed two co-culture models wherein organoids can be cultured with endothelial cells that form a micro-vessel network. We found that intestinal organoids and micro-vessels have different medium requirements. Intestinal organoids impair micro-vessel network formation, while endothelial cells can increase stem cell maintenance in organoids. Our work can form the basis for future studies on vascularization of organoids *in vitro*.

## Results

### Generating a microvessel network in a microfluidic device

Microfluidic devices are powerful tools for facilitating novel models for complex cocultures. To explore the possibilities of using the AIM Biotech DAX1 microfluidic devices (chips) for studying organ and tumor vessels we used various protocols to co-culture organoids and vessels in these chips. We first established a simple protocol in which a vasculature develops through the process of vasculogenesis in the chips. In this procedure Human Umbilical Vein Endothelial Cells (HUVECs) were co-cultured in the center channel of the chips in a fibrin matrix with VH10 fibroblasts (**Figure 1A**). The side-channels were filled with cell culture medium (EGM2). The HUVECs are initially visible as trypsinized single cells on day 1.These cells become elongated sprouts on the second day, which become more connected by day 4/5. Following this the endothelial cells start proliferating to form a mature network of microvessels with a maximum length at Day 5-7. (**Figure 1B-C**). The density and lumenization of the endothelial network depends on the ratio of the cell types used; we found that using an equal number of HUVECs vs Fibroblasts or a larger number of HUVECS yields better connected vascular networks (**Figure 1D-E**). Indeed, by confocal imaging we found that using high cell numbers generates microvessels that are lumenized (**Figure 1F**).

**Figure 1.**
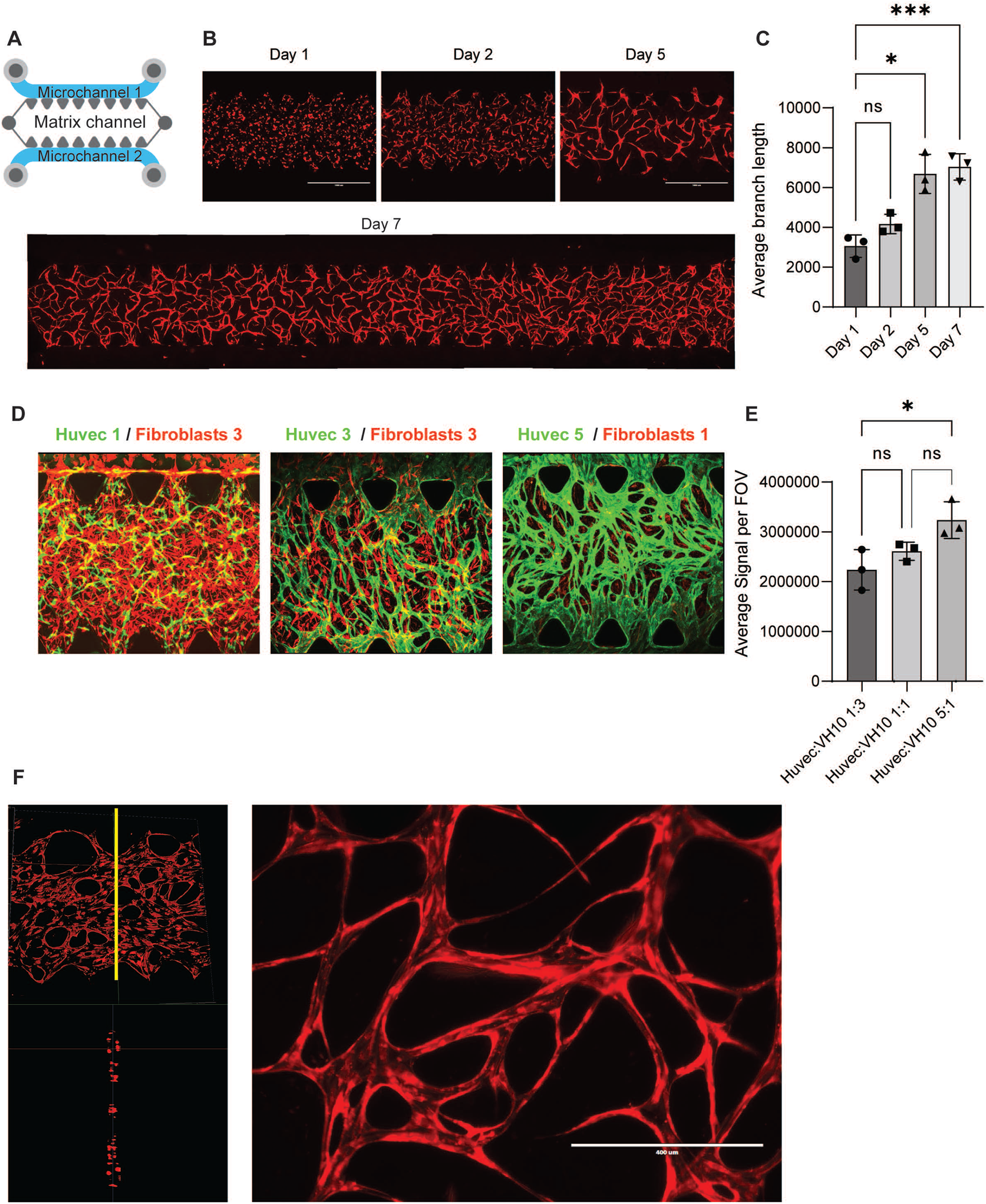
Generating a microvessel network in a microfluidic device. **(A)** Schematic representation of AIM Biotech microfluidic device (chip) containing 3 channels. The middle channel is used for matrix (with cells), while the outer channels are used for medium. **(B)** Representative EVOS microscope images of HUVEC_DsRed in a vasculogenesis assay. HUVEC_DsRed and fibroblasts were cultured in a fibrin matrix in the center channel of the chip; the side-channels contain EGM2. HUVEC_DsRed form a remodelling vascular network over the course of a week. **(C)** Quantification of average branch length of vascular network by FIJI.Data is represented as average relative branch length +/-standard deviation (SD). P-values were calculated by ANOVA using Dunnett test for multiple comparisons (*p<0.05, ***p<0.001) **(D)** Representative confocal images of HUVEC_GFP and Fibroblasts_DsRed in a vasculogenesis assay on day 7. Images are maximum-projections of z-stacks. Concentrations used were 1 and 3 million/ml, 3 and 3 million/ml or 5/1 million/ml (Huvec and fibroblasts respectively). **(E)** Coverage per field-of-view refers to quantification of GFP-signal per field of view and gives a quantitative measure of the size of the HUVEC network. Data is represented as average -/+ SD. P-values were calculated by ANOVA using Dunnett test for multiple comparisons (*p<0.05) **(F)** Representative confocal images of HUVEC_DsRed in a vasculogenesis assay on day 7. Images are maximum-projections of z-stacks. Image on the left shows cross-sections of z-stacks indicating lumenization of vascular network.

### Co-culture of organoids, EC’s and fibroblasts in a vasculogenesis assay

Next, we addressed whether these microvessels can be co-cultured with organoids. Human Small Intestinal (hSI) organoids are a well-characterized organoid model and we therefore used these to develop a co-culture model in the AIM Biotech chip. The challenge for such a co-culture model lies in the different conditions that the cells are cultured in. Organoids are generally cultured in matrigel in organoid medium, while the microvessels perform optimally in fibrin in EGM2 we first aimed to assess whether we could define a matrix that would allow culturing of both of these cell types. As it has been reported that human SI organoids can grow in fibrin/matrigel hybrid matrices ^34^ we set out to assess whether a fibrin/matrigel can serve as a matrix to allow culturing of hSI organoids. While we found that hSI organoids cannot be cultured in a 100% fibrin matrix (data not shown), they do grow well in mixtures containing 10 or 20% of matrigel (**Supplementary Figure 1A-B**) in organoid medium. Next, we also addressed whether various mixtures of organoid medium and EGM2 were able to sustain organoid growth. Our data showed that mixtures with EGM2 are remarkably well able to support organoid growth in short cultures. After four days of culture various mixtures of EGM2 and organoid medium, ranging from 1:5 to 5:1, all were able to allow organoids to grow and survive (**Suppelementary Figure 1C-D**).

We also assessed whether addition of matrigel to the fibrin matrix was compatible with microvessel formation in the chip. These results showed that addition of 10% matrigel did not affect vessel formation while 20% of matrigel resulted in a small reduction of branch formation (**Supplementary Figure 2E-F**). In addition, by testing out a variety of medium compositions we determined that a composite medium of 2:1 (EGM2:hSI EM) was optimal for network formation in the fibrin/10%matrigel matrix (**Supplementary Figure 2G-H**). Since these conditions also seemed favorable to organoid growth we setup triple co-cultures (organoids, fibroblasts and endothelial cells) in the chips.

After having defined optimal conditions to culture microvessels and organoids separately, we combined these conditions for a triple culture of HUVECs, fibroblasts and hSI organoids in a fibrin/MG10% matrix in composite (2:1 EGM2:organoid) medium. We compared the triple culture also to mono-culture of organoids or to co-culture of HUVECs and fibroblasts. We found that hSI organoids do not survive well in mono-culture (**Figure 2A**). On day 2, the majority of the organoids in mono-culture have become dense, indicating (partial) differentiation (**Figure 2A-B**). In contrast, in combination with HUVECs and fibroblasts the organoids maintain their cystic nature better, which is a measure for remaining in stem-like state (**Figure 2A-B**). However, after 4 days most organoids become differentiated and die also in the triple culture (**Figure 2C**). To improve the survival of organoids in our model we hypothesized that increasing Wnt signaling in the cultures may be relevant as Wnt signaling is a crucial determinant in effecting stem cell maintenance and self-renewal in the intestinal stem cells. We set up triple cultures in various concentrations of CHIR 99021 (CHIR), which inhibits glycogen synthase kinase (GSK) 3β, a key negative regulator of Wnt signaling. We observed that indeed addition of CHIR leads to a higher number of cystic organoids indicating that an increase in Wnt signaling can improve survival of organoids in our co-culture model (**Figure 2D-E**). We found that addition of 5 nM CHIR yields a high percentage (of 73%) of cystic organoids by day 4 compared to only 13% without CHIR **(2E)**. Additionally, the number of formed junctions and branches by HUVECs significantly increased with 1nM of CHIR and remained high in 5nM CHIR **(Figure 2E)**. These findings indicate that triple-cultures require additional Wnt-stimulation to allow organoids to maintain stemness in the co-cultures in the chips.

**Figure 2.**
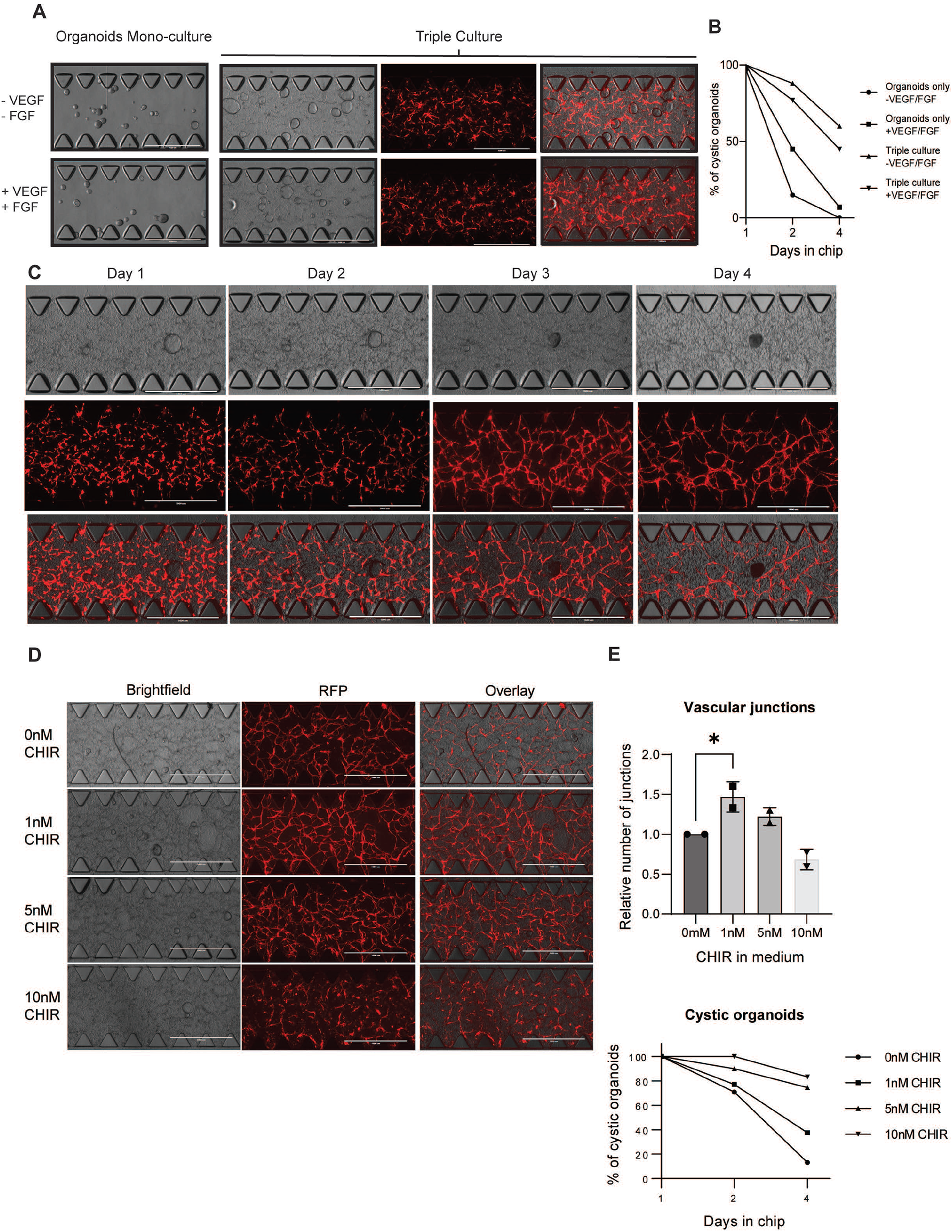
Co-culture of organoids, EC’s and fibroblasts in a vasculogenesis assay. **(A)** Representative EVOS images of organoids and HUVEC_DsRed cultured alone or in a triple co-culture (organoids, HUVEC_DsRed and fibroblasts) in a vasculogenesis assay in the microfluidic chip. **(B)** Quantification of cystic organoids in triple co-culture with fibroblasts and HUVEC or cultured as mono-culture with or without VEGF and FGF. Data is represented as percentage of all quantified organoids. **(C)** Representative EVOS images of organoids and HUVEC_DsRed cultured in a triple co-culture (organoids, HUVEC_DsRed and fibroblasts) in a vasculogenesis assay in the microfluidic chip. HUVEC_DsRed form a vascular network over time. Organoids shrink and become dense by day 3 and are mostly dead (very dark) by day 4. **(D)** Representative EVOS images of organoids and HUVEC_DsRed cultured in a triple co-culture (organoids, HUVEC_DsRed and fibroblasts) in a vasculogenesis assay in the microfluidic chip in conditions that vary in the added CHIR. **(E)** Quantification of total number of branches of vascular network and average organoid size by FIJI. Data is represented as mean +/-standard deviation (SD). P-values were calculated by ANOVA using Dunnett test for multiple comparisons (*p<0.05) **(F)** Quantification of cystic organoids in triple co-culture with fibroblasts and HUVEC in different concentrations of CHIR. Data is represented as percentage of all quantified organoids.

### Direct cell-to-cell contact between ECs and Fibroblasts is not required for establishing a microvessel network

To explore whether the endothelial cells require direct cell-to-cell contact we also tested the possibility of culturing the fibroblasts in the side-channels of the chips. We found that HUVECs can readily form endothelial networks using this setup, indicating that paracrine signaling by fibroblasts is sufficient to stimulate vasculogenesis (Data not shown). We also showed that similar vascular networks could be generated using the Endothelial Colony Forming Cells that constitutively express GFP (ECFC_GFP) (**Figure 3A-B**). Similar to HUVECs these endothelial cells form networks which can remodel into lumenized microvessels over time (**Figure 3C**). We analyzed these networks by confocal microscopy and we found that these microvessels are hollow lumenized structures by generating cross sections (**Figure 3D**). To determine whether these vessels do form a functional, connected network we injected mCherry labeled MDA-MB-231 cells, a breast cancer cell line, into the side channels of the chip. By confocal imaging we found that the MDA-MB-231 cells were transported through the vessels and not through the fibrin matrix, indicating that these microvessels are functional and non-leaky (**Figure 3E**). These data indicate that direct cell-to-cell contact between ECs and Fibroblasts is not required for establishing a microvessel network.

**Figure 3.**
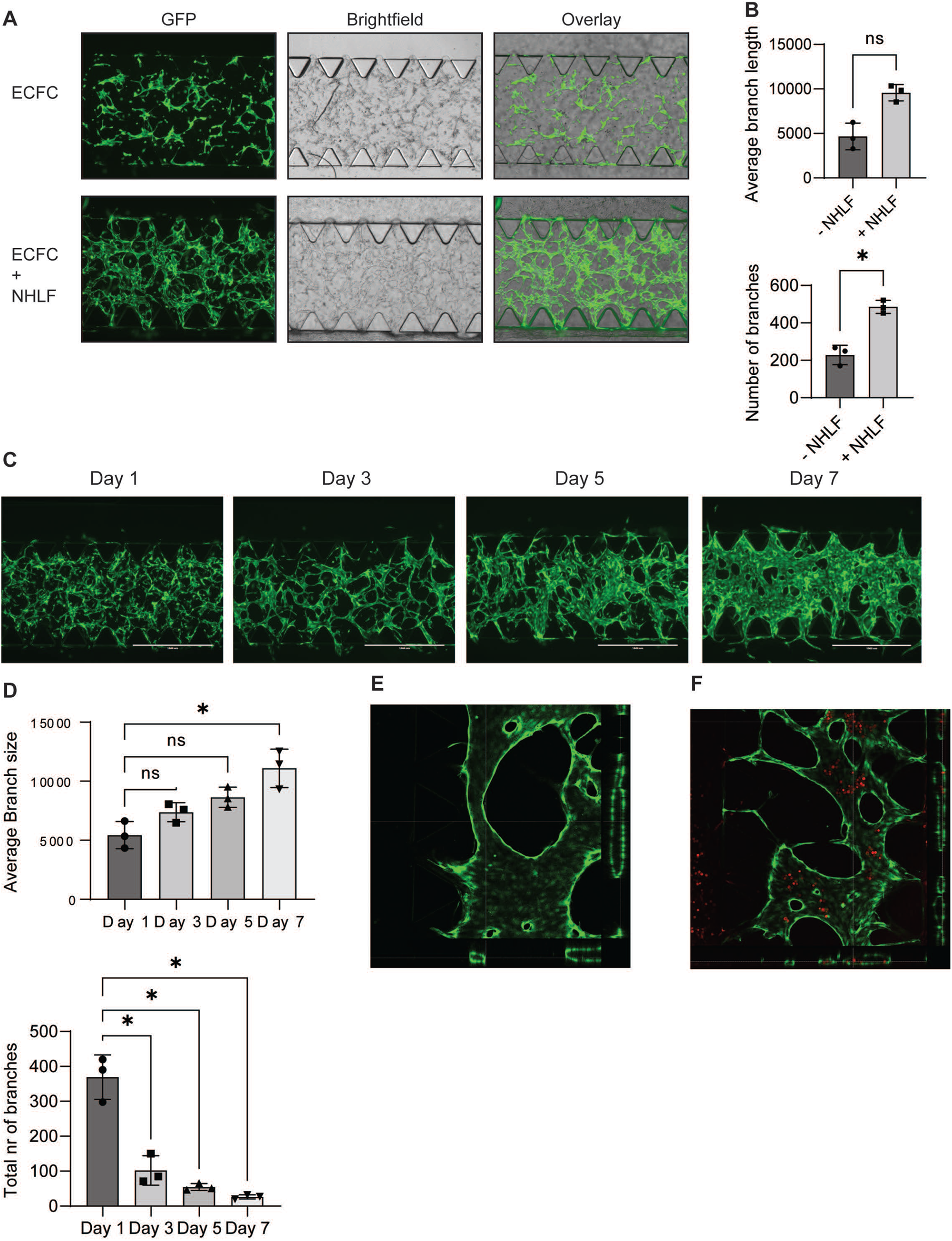
Direct cell-to-cell contact between ECs and Fibroblasts is not required for establishing a microvessel network. **(A)** Second method wherein ECFC were cultured in a fibrin matrix in the center channel of the chip. Images are representative images of ECFC_GFP and brightfield (to show non-labeled fibroblasts). Experiments were setup without (upper image) and with fibroblasts (NHLF, bottom images) in the side-channels. **(B)** Quantification of average branch length and number of branches of vascular network of ECFC_GFP by FIJI. Data is represented as mean +/-standard deviation (SD). P-values were calculated by paired t-test (ns p>0.05, *p<0.05) **(C)** Representative EVOS microscope images of ECFC_GFP vascular network. ECFC_GFP were cultured in fibrin matrix with fibroblasts in side-channels in EGM2. ECFC_GFP form a remodelling vascular network over a timecourse of a week. **(D)** Quantification of average branch size and number of branches of vascular network of ECFC_GFP by FIJI. Data is represented as mean +/-standard deviation (SD). P-values were calculated by paired t-test (ns p>0.05, *p<0.05). In this network the number of branches goes down, while the branch size goes up indicating that vessels become larger and more connected to each other. **(E)** Representative confocal images of ECFC_GFP in second vasculogenesis assay on day 7. Images are maximum-projections of z-stacks. On the bottom and right cross-sections of z-stacks are shown indicating lumenization of vascular network. **(F)** Representative confocal images of MDA_MB_231_RFP and ECFC_GFP in second vasculogenesis assay on day 7. MDA_MB_231_RFP were injected as single cells into the side-channels and were found to travel through the vascular network. Images are maximum-projections of z-stacks.

### Co-culturing organoids and ECFC in a separate compartment from HUVECs leads to different responses to medium conditions

We then explored how organoids can be cultured in this setup. Hence, the organoids and endothelial cells were co-cultured in a fibrin/matrigel (90/10) matrix in the center channel of the chip, while fibroblasts were cultured in the sidechannels (**Figure 4A**). We tested various medium conditions and assessed the microvessel network development by quantifying the branch length of the microvessels (**Figure 4B**). In this setup we found that co-culturing organoids together with endothelial cells impairs the development of the microvessel network even in pure EGM2 (**Figure 4A-B**). We also tested adding CHIR in this setup, but while this does lead to better organoid growth, this also induces a stronger impairment of endothelial network formation in this method (**Figure 4A-B**). In agreement, using the composite medium of 2:1 (EGM2:hSI-EM) that has a higher proportion of organoid medium, had a positive effect on organoid size, but the microvessel network was less branched and showed less mature vessels (**Figure 4A-B**). These findings show that the effects of medium composition can have different effects depending on the experimental setup.

**Figure 4.**
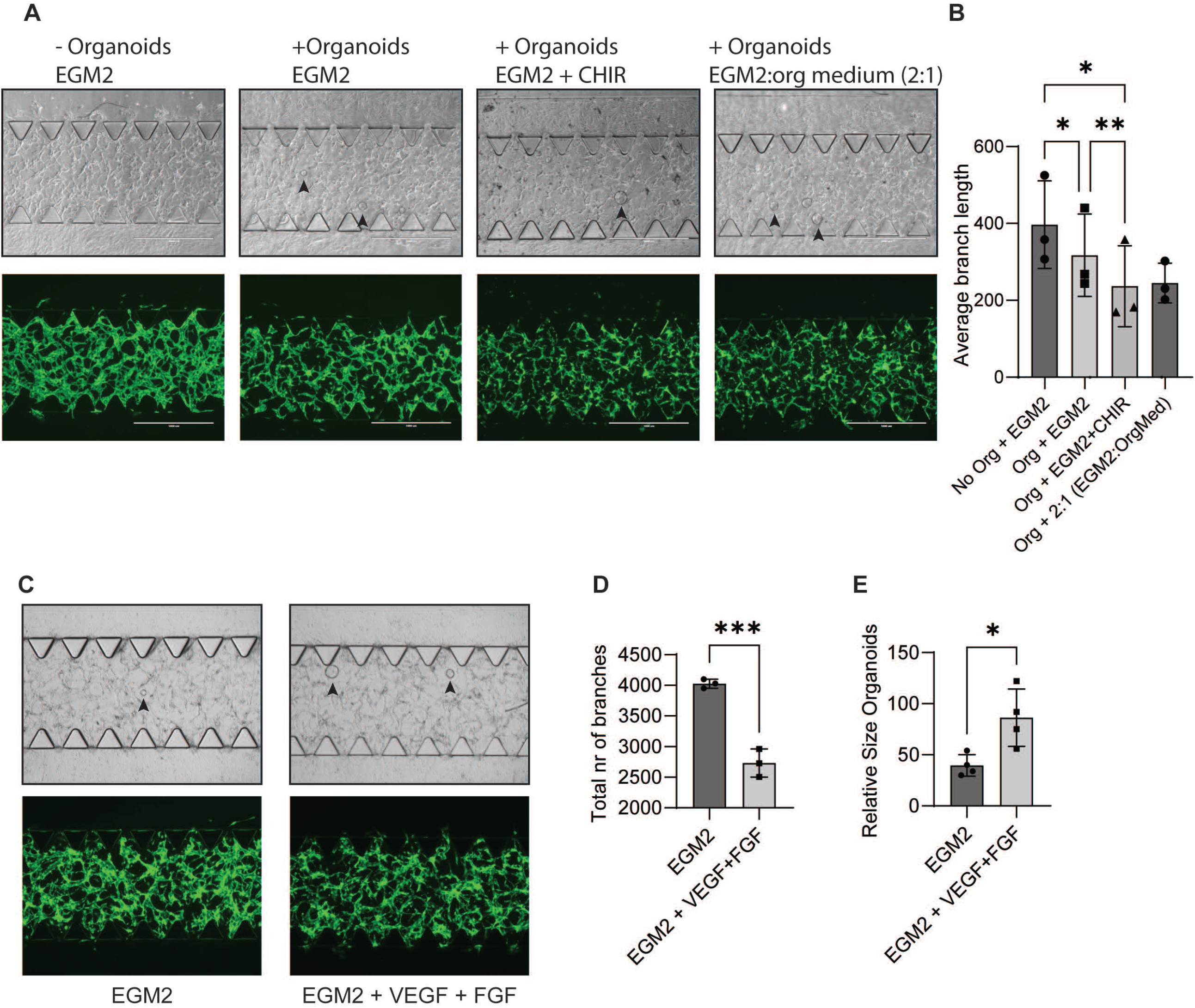
Co-culturing organoids and ECFC in a separate compartment from HUVECs leads to different responses to medium conditions. **(A)** Representative EVOS images of ECFC_GFP vascular network and brightfield images to identify organoids. ECFC_GFP were cultured in a fibrin/matrigel matrix in the center channel without/with organoids. Side-channels were filled with fibroblasts and various medium compositions as indicated. Arrowheads highlight organoids. **(B)** Quantification of average branch length of vascular network by FIJI. Data is represented as average relative branch length +/-standard deviation (SD). P-values were calculated by ANOVA using Dunnett test for multiple comparisons (*p<0.05, **p<0.01) **(C)** Representative EVOS images of ECFC_GFP vascular network and brightfield images to identify organoids. ECFC_GFP were cultured in a fibrin/matrigel matrix in the center channel without/with organoids. Side-channels were filled with fibroblasts in EGM2 with or without VEGF (50ng/ml) and FGF (50ng/ml). Arrowheads highlight organoids. **(D)** Quantification of total number of branches of vascular network and average organoid size by FIJI. Data is represented as mean +/-standard deviation (SD). P-values were calculated by ANOVA using Dunnett test for multiple comparisons (***p<0.001,*p<0.05)

To further improve the endothelial network development in the presence of intestinal organoids we explored how the addition of angiogenic growth factors affected the co-cultures. We chose to use Vascular Endothelial Growth Factor (VEGF) and Fibroblast Growth Factor (FGF) as these are well-characterized to stimulate angiogenesis in vivo and in vitro (REF). Under these experimental conditions, the addition of VEGF and FGF does not lead to a more mature endothelial network and even yields a lower number of branches (**Figure 4C-D**). Interestingly, addition of VEGF, FGF or the combination did positively affect the organoid size (**Figure 4C,4E**). In addition we found that in EGM2 a large majority of organoids acquire a dense differentiated phenotype (**Figure 4C,4E**). In the presence of VEGF and/or FGF, a higher proportion of the organoids remained cystic in comparison to co-culture in EGM2 (**Figure 4D**). Together these data show that addition of FGF and VEGF leads to a more sprout-like endothelial network while organoids are more prevalently found in a stem-like state.

### Angiocrine signaling as a possible mechanism for mediating organoid stemness

From the previous experiments we were unable to distinguish whether FGF and VEGF had a direct effect on organoid stemness or whether this required the presence of microvessels. To test this we compared organoids in monoculture in organoid medium, in monoculture in EGM2 (+ VEGF and FGF) and in co-culture in EGM2 (+ VEGF and FGF). We found that organoids in EGM2 +VEGF + FGF were unable to sustain their normal cystic phenotype. After 24 hours most of the organoids are unable to grow or they become cystic (**Figure 5A**). Unexpectedly, even in full organoid medium, the organoids are largely unable to grow in the chip (**Figure 5B**). This perhaps reflects the inability of the medium to travel through the matrix. However, in the co-cultures, the organoids were able to grow and they remained cystic for several days (**Figure 5C**). Quantification shows that organoids in co-culture with ECFC/NHLF are larger and that a higher proportion of the organoids are cystic for a longer period of time (**Figure 5C**). These data suggest that angiocrine signaling from the ECFC/NHLF induces stem cell maintenance in organoids.

**Figure 5.**
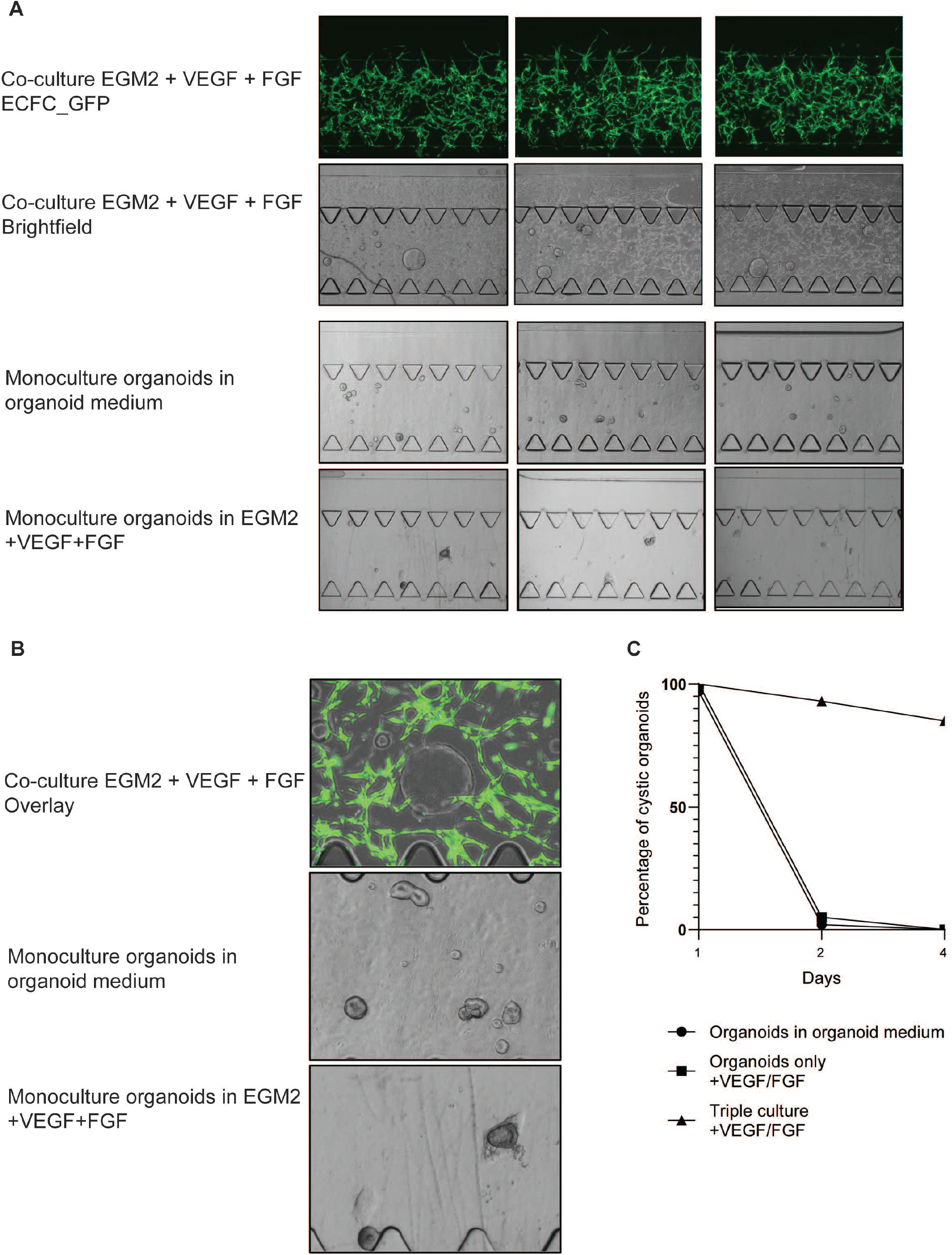
Angiocrine signaling as a possible mechanism for mediating organoid stemness. **(A)** Representative EVOS images of ECFC_GFP vascular network and brightfield images to identify organoids. Organoids were cultured in a fibrin/matrigel matrix in the center channel with/without ECFC_GFP. Side-channels were filled with fibroblasts and various medium compositions as indicated. **(B)** Representative zoom in of EVOS images of ECFC_GFP vascular network and brightfield images to identify organoids of the same experiments.

## Discussion

In this manuscript, we discuss our work on developing models to study interactions between vasculature and organoids. We envision that these models could be used to study organ vascular development or diseases that affect multiple cell types, such as inflammatory bowel diseases, or cancer. In addition, a better understanding of how to co-culture blood vessels with organoids could lead to developing vascularized organoids and form a strong base for using organoids for regenerative medicine purposes.

We established two different protocols to use the chips to study organ vascularization. In the first method, the fibroblasts were cultured inside the same fibrin/matrigel matrix as the organoids and ECs. Here, EC’s self-organize into a vascular network, but we found that co-culturing of organoids represents with a number of technical challenges. First, in the chip organoids exhibit a limited lifespan, which is most prominently apparent in mono-culture where virtually all organoids die after 4 days (**Figure 2C**). This perhaps reflects an inability to access the full proportion of signaling factors (with Wnt signaling being the most important affected pathway) in the medium. Moreover, the matrix in the chip is in a confined space, which does not allow the matrix to expand in size, and this will most likely also increase the pressure on the organoids. In the triple-culture setup this can be mitigated as organoids exhibit a higher proportion of cystic (stem-like) phenotype. Enhancing Wnt-signaling by CHIR can partially rescue these impairments and seems like the best way of allowing organoids to grow together with a developing vascular network.

In contrast, in our second method, where the fibroblasts are cultured in separate channels the addition of CHIR did not result in a mutual beneficial condition. Under these experimental conditions we found that conditions that favoured vessel formation also were optimal for co-culture of organoids and microvessels. Thus, the setup of the co-culture determines that the cells can respond differently to changes in experimental conditions. This illustrates the complexity of these co-culture systems and highlights the context-dependent responses in biological systems. The second method overall has the best possibility of retaining organoids in a stem-like growing state, while microvessels form a connected vasculature.

Interestingly, organoids and microvessels have complex effects on each other in the chip. We found that co-culture of micro-vessels can stimulate organoid stemness and survival. This is especially prominent in the second method where the organoids are even able to be cultured in pure EGM2 (+ VEGF/FGF). This most likely is an effect of angiocrine signaling from the endothelium to the organoids. We propose that RSPO3 is a good candidate to mediate these effects. RSPO3 has been reported to be expressed in the intestinal niche ^35,36^ and we found it to be the only secreted modulator of Wnt signaling to be expressed in HUVECs and ECFC (data not shown). Moreover, the micro-vessels may affect the matrix stiffness which is also a crucial factor in maintaining organoid stemness ^37,38^.

On the other hand, the organoids have an inhibitory effect on vascular network formation. We believe that this is a result from the organoids competing for nutrients, metabolites and oxygen. ECs are known for their high metabolic consumption rate during angiogenesis^39–41^ and this may be hampered by organoids. In addition the organoids also affect matrix remodeling, perhaps physically obstructing the way that endothelial cells can migrate. We also found that organoids exhibit very low expression of angiogenic factors such as VEGF and FGF (data not shown). Perhaps tellingly, blood vessels in the intestine do not necessarily are in close contact with the epithelium, but rather remain at a (small) distance ^42^. *In vivo* often other cell types, such as macrophages, astrocytes, fibroblasts, smooth muscle cells have been reported to direct angiogenesis and perhaps the lack of these cells would have to be compensated. The difficulty of using ASC-derived organoids for these models is that each required cell type will have to be added individually. Perhaps, using IPSC-derived organoids can allow for different cell types to be incorporated from the same source.

Together, we have developed co-culture assay that allows organoids to be co-cultured with a microvessel network. These cells affect each other by competing for nutrients, affecting their extracellular environment and by signaling to each other. This study can form the basis for future work to develop a fully functional vasculature in organoid cultures.

## Materials and Methods

### Cell culture

#### HUVECs and ECFC

ECFC isolation and culture has been described previously Human umbilical endothelial cells (HUVECs) from Lonza were cultured in T-75 cell culture flasks in Endothelial Cell Growth Medium-2 (EGM-2) (BulletKit, Lonza, Cat. No. CC-3162). EGM2 was made from Endothelial cell Basal Medium – 2 (EBM2), with added endothelial supplements including 2% fetal bovine serum (FBS) (v/v), hydrocortisone, VEGF, human FGF, R3-IGF-1, ascorbic acid, human EGF, glutaraldehyde GA-1000, and heparin. The medium was refreshed 3 times per week, and cells were passaged (p) by trypsinization weekly at ca. 70% confluency. After trypsinization, cells were seeded at 2×10^5^ cells per new flask. HUVECs were taken into culture at p3.

#### VH10

VH10 foreskin fibroblasts (kind gift from Ag Jochemsen, LUMC) were cultured in Dulbecco’s Modified Eagle Medium (DMEM) with high glucose, sodium pyruvate and GlutaMAX-1 (Gibco, Cat. No. 10569010) supplemented with 10% FBS (v/v) and 1% penicillinstreptomycin (P/S) (v/v) (Gibco, Cat. No. 15140122).

#### NHLF

NHLF (Normal Human Lung Fibroblasts, Lonza) were cultured in Fibroblast Growth Medium 2 (FGM2), with added supplements. The medium was refreshed 3 times per week, and cells were passaged (p) by trypsinization weekly at ca. 70% confluency. After trypsinization, cells were seeded at 5×10^5^ cells per new flask.

#### HEK

Transformed human embryonic kidney cells HEK293T (kind gift from Martijn Rabelink, LUMC) were cultured in DMEM with high glucose, sodium pyruvate and GlutaMAX-1 supplemented with 10% FBS (v/v) and 1% P/S (v/v) in 10cm cell culture plates.

#### Human small intestinal organoids

Human small intestinal cell organoids (hSIOs) were obtained from healthy donor duodenum biopsies. All participants gave written consent for participation in this study, following a protocol approved by the review board of the UMC Utrecht, The Netherlands (protocol STEM study, METC 10-402/K).

Organoids were grown in Matrigel diluted with 30% DMEM/F-12 (Gibco, Cat. No. 11320033) supplemented with 1% HEPES (v/v), 1% P/S (v/v) (Gibco, Cat. No. 15140122). 3 droplets of 10ul were plated per well of a 24-well suspension culture plate. Human intestinal stem cell expansion medium (hSI EM) was used to culture the organoids. Organoids were either collected for experiments at 3-5 days post-passage or passaged again after a total 5-8 days of culture. Organoids were passaged either into single cells by TryplE express enzyme (Gibco, Cat. No. 12604013), and 2×10^4^ single cells were plated per new well. After passaging by single cells, 1ul/ml of rock inhibitor Y27632 was added to the medium for the first 48h post-passage. Alternatively organoids were physically disrupted into small fragments by pushing organoids 10-15 times through a 200ul pipette tip, and organoid fragments were split in a 1:4 ratio into new wells.

#### Generation of HUVEC_DsRed

Lentiviral constructs for transduction of cells were packaged using HEK293T cells. On day 0, 4.5×10^5^ cells were seeded on ø10cm cell culture plates and cultured in DMEM with high glucose, sodium pyruvate and GlutaMAX-1 (Gibco, Cat. No. 10569010) supplemented with 10% FBS (v/v) and 1% P/S (v/v) (Gibco, Cat. No. 15140122).

On day 1, cells were transfected with 10ug of one of the lentiviral constructs carrying the gene of interest and two packaging plasmids, 10ug PAX2 (Addgene, Cat. No. 12260) and 5ug VSV.G (Addgene, Cat. No. 14888).

The construct of interest and packaging plasmids were incubated in serum-free medium with 1:5 PEI reagent for 30min and then added dropwise to the cells. Transfected cells were left to incubate overnight and refreshed on day 2. On day 3, the virus-containing supernatant was harvested and refreshed with new medium, which was collected in a second harvest on day 4. Virus-containing medium was centrifuged for 4min at 1600rpm. Any debris was discarded, and the supernatant was filtered through a 45µm syringe filter, collected in a new tube, and stored at -80°C until use.

To generate HUVECs overexpressing DsRed unlabelled HUVECs (p0) were seeded at 1×10^5^ cells per well in a 6 well cell culture plate. Each well was supplied with 1.5ml of EGM2. Cells were allowed 1-2 days to attach, before they were transduced. Near-confluent cells were transduced by aspirating the medium and adding 1ml of the respective virus diluted in 1ml of EGM2 with 4mg/ml polybrene (Sigma, Prod. No. H9268-5G). Cells were then left to incubate for 6h at 37°C and 5% CO_2_. After 6h, the virus was aspirated and cells were refreshed with new medium. HUVEC were found to be >90% positive for DsRed using this method and were used without further enrichment of positive cells.

#### Set-up of vasculogenesis assay with triple culture in matrix channel in the microfluidic device

The matrix holding cells in the vasculogenesis assay was fibrin matrix with 10% Matrigel. Fibrinogen from bovine plasma (Sigma, Cat. No. F8630-1G) was dissolved in PBS supplemented with *Ca*2+ and *Mg*2+ ions at a concentration of 5mg/ml. After the solution was homogenous, it was filtered through a 0.2µm syringe filter. The solution was then later used to resuspend the harvested cells.

2-3 wells of organoids were collected 4-5 days post-passage from culture plates by dissolving the Matrigel droplets in ice-cold GF-. Organoids were spun at 800rpm for 3min. The supernatant was removed and the remaining organoids were re-dissolved in ice-cold GF-. VH10 and HUVECs were harvested by trypsinization at ca. 90-100% respectively 70% confluency. VH10 and HUVECs were centrifuged at 1500rpm for 4 minutes to generate a pellet. Supernatant was aspirated and the pellet resuspended in culture medium. 6×10^5^ VH10 and 6×10^5^ HUVECs (respectively 2×10^5^ and 10×10^5^ for respective experiments) were mixed with 2-3 wells of organoids, and centrifuged at 1100rpm for 5 minutes to generate a pellet. Supernatant was aspirated. Cells were dissolved in 90µl of 5mg/ml fibrinogen solution which was then supplemented with 10µl Matrigel, for a final concentration of 6×10^6^ cells/ml (respectively 2×10^6^ and 10×10^6^ cells/ml for respective experiments) of each HUVEC and VH10. To 100µl of cell-suspension, 3µl of thrombin were added and quickly mixed by pipetting. Immediately after, to avoid polymerization outside of the microfluidic device, 10µl of cell-suspension were injected into the middle channel of a microfluidic site. The device was left to incubate for 10min at room temperature followed by 20min at 37°C and 5% CO_2_.

After incubation, the media channels were filled with 15µl of 2:1 EGM2:hSI EM medium (EOM), organoid medium, or organoid medium supplemented with the endothelial supplements of EGM2, including 2% fetal bovine serum (FBS) (v/v), hydrocortisone, VEGF, human FGF, R3-IGF-1, ascorbic acid, human EGF, glutaraldehyde GA-1000, and heparin. For experiments with PI20-transduced HUVECs, 1µg/ml doxycycline was added to the medium. 70µl or 50µl of the medium were filled in opposite media ports of each channel. The water reservoirs in the chip holder plate were filled with MilliQ water and the microfluidic device was placed in the incubator at 37°C and 5%CO_2_. The triple co-cultures were maintained for 5 days with daily media refreshments.

#### Set-up of vasculogenesis assay with duo culture in matrix channel in the microfluidic device with fibroblasts in side-channels

The matrix used was prepared similarly as described for the triple culture. In addition, the organoids, were prepared as above, while ECFC were prepared like HUVECs in a similar procedure. 6×10^5^ HUVECs (respectively 2×10^5^ and 10×10^5^ for respective experiments) were mixed with 2-3 wells of organoids, and centrifuged at 1100rpm for 5 minutes to generate a pellet. Supernatant was aspirated. Cells were dissolved in 90µl of 5mg/ml fibrinogen solution which was then supplemented with 10µl Matrigel, for a final concentration of 6×10^6^ cells/ml of HUVEC. To 100µl of cell-suspension, 3µl of thrombin was added and quickly mixed by pipetting. Immediately after, to avoid polymerization outside of the microfluidic device, 10µl of cell-suspension was injected into the middle channel of a microfluidic site. The device was left to incubate for 10min at room temperature followed by 20min at 37°C and 5% CO2.

After incubation, the media channels were filled with 70 ul and 50 ul of a suspension of NHLF or VH10 (2 million/ml).

#### Microscopy

Vasculogenesis assays in the microfluidic device were imaged by an EVOS fluorescent microscope using a 4x objective. Brightfield, RFP/GFP, and overlay images were acquired.Confocal microscopy was performed using a Leica SP8 confocal microscope.

#### Fiji - ImageJ analyses of vascular networks

To assess network formation by HUVECs in the vascularization assay in the microfluidic device, fluorescent and confocal microscopy images of networks on day 4 were analysed in Fiji (ImageJ). Images were one by one loaded into Fiji and colour threshold was manually adjusted by navigating ‘Image’->‘Adjust’->‘Color threshold’ to exclude background noise as much as possible. Next, the image was made binary by the ‘Process’->‘Binary’->‘Make binary’ command. The image was then smoothened by applying a Gaussian Blur at ‘Process’->‘Filters’->‘Gaussian Blur’ (radius 2.00), and made binary again following the described process. The binary image was skeletonized (‘Process’->‘Binary’->‘Skeletonize’) and the resulting skeleton was analysed by navigating the ‘Analyze’->‘Skeleton’->‘Analyse skeleton’ commands. From the results table, the columns for number of branches and number of junctions were imported to Microsoft Excel, where the sum was calculated for each variable. For multiple pictures of the same channel, the sums of branches and junctions were averaged. For repeated experiments, replicates within each experiment were averaged. The number of branches and junctions of each condition was normalized by the control condition and plotted in GraphPad Prism software.

#### Organoid quantification

To assess survival of organoids in the microfluidic device, organoids were classified as either cystic, dense, or apoptotic at 1, 2, and 4 days after introduction into the microfluidic device. Classification was done manually based on brightfield microscopic images. Cystic organoids were defined as round and transparent, without signs of differentiation or apoptosis. Organoids with a darkened colour but still a defined shape were classified as ‘dense’. Apoptotic organoids were defined by a dark colour and loss of a defined shape, showing signs of disintegration and shredding of single cells or apoptotic bodies. Counts for each phenotype were collected in Microsoft Excel, where the percentage of each phenotype of the total number of organoids was calculated.

## Acknowledgments

This collaboration project is co-funded by the PPP Allowance made available by Health∼Holland, Top Sector Life Sciences &Health, to stimulate public-private partnerships. This project was further supported by funding by Stichting Proefdiervrij and the 3V Stimuleringfonds Utrecht. We are grateful to Kuan Chee Mun, Sei Hien Lim and Lawrence Lim, Cindy Frederiks, Iris Schilt, Josse Bouwhuis, Jennifer Sarikaya for advice and help during the project.

## Figure Legends

**Supplementary Figure 1.**
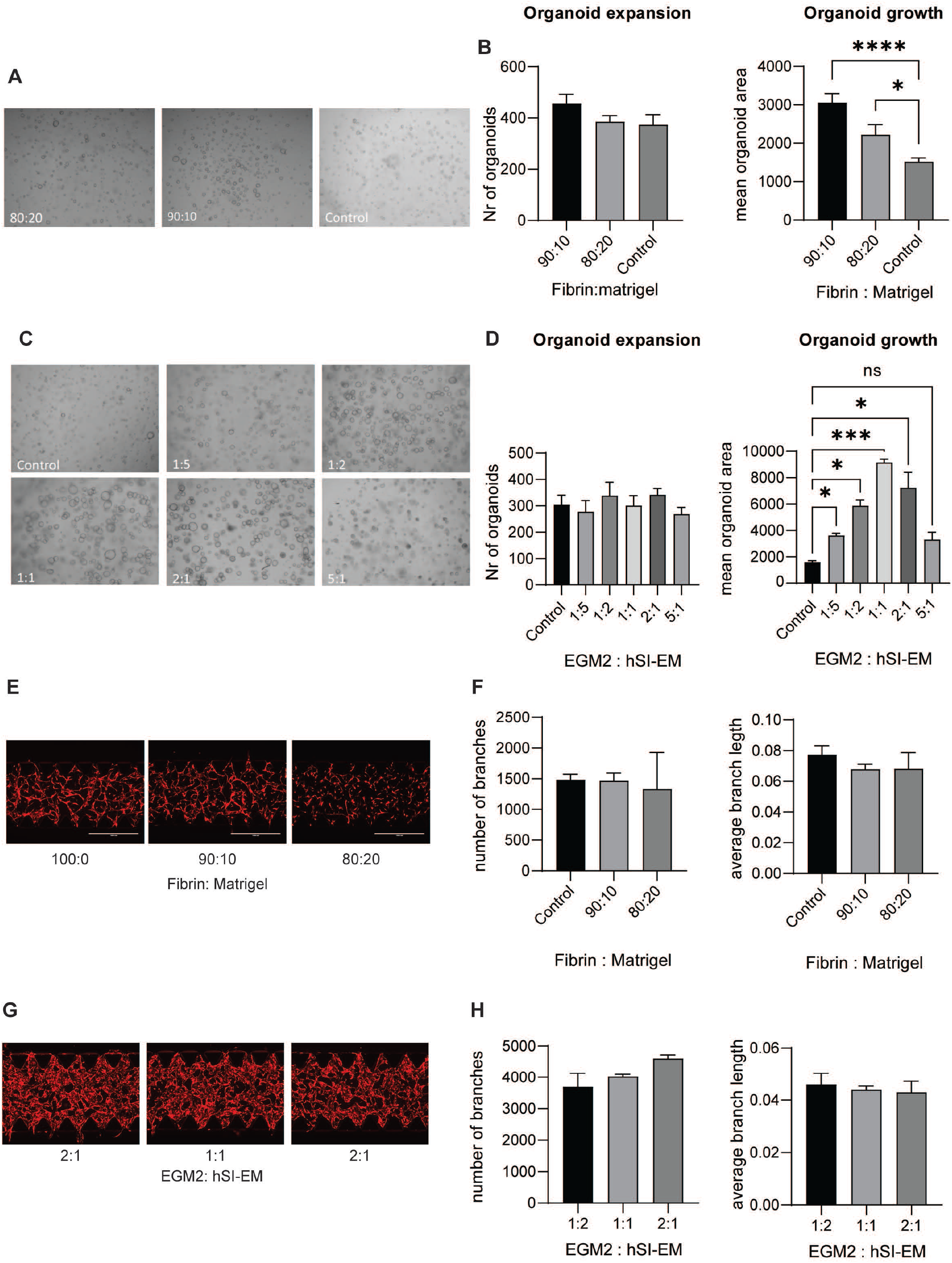
Optimizing medium conditions for organoids and micro-vessels. **(A)** Representative EVOS images of organoids growing in different matrix compositions. **(B)** Left:Quantification of total number of organoids. N=3, a minimum of 3 images per experiment were counted. Data is represented as mean +/-SD. P-values were calculated by ANOVA using Dunnett test for multiple comparisons (all non significant) Right: Quantification of organoid size. N=3, a minimum of 3 images per experiment were counted. Data is represented as mean +/-SD. P-values were calculated by ANOVA using Dunnett test for multiple comparisons (****p<0.0001,*p<0.05) **(C)** Representative EVOS images of organoids growing in different medium compositions. **(D)** Left:Quantification of total number of organoids. N=3, a minimum of 3 images per experiment were counted. Data is represented as mean +/-SD. P-values were calculated by ANOVA using Dunnett test for multiple comparisons (all non significant) Right: Quantification of organoid size. N=3, a minimum of 3 images per experiment were counted. Data is represented as mean +/-SD. P-values were calculated by ANOVA using Dunnett test for multiple comparisons (***p<0.001,*p<0.05) **(E)** Representative EVOS images of HUVEC_DsRed cultured in co-culture with fibroblasts in a vasculogenesis assay in the microfluidic chip in different matrix compostions. **(F)** Quantification of average branch length and number of branches of vascular network of HUVEC_DsRed by FIJI. Data is represented as mean +/-standard deviation (SD). P-values were calculated ANOVA using Dunnett test for multiple comparisons (all non significant). **(G)** Representative EVOS images of HUVEC_DsRed cultured in co-culture with fibroblasts in a vasculogenesis assay in the microfluidic chip in different medium compostions. **(H)** Quantification of average branch length and number of branches of vascular network of HUVEC_DsRed by FIJI. Data is represented as mean +/-standard deviation (SD). P-values were calculated ANOVA using Dunnett test for multiple comparisons (all non significant).

